# Heterogeneity analysis of acute exacerbations of chronic obstructive pulmonary disease and a deep learning framework with weak supervision and privacy protection

**DOI:** 10.1101/2023.12.04.570028

**Authors:** Yuto Suzuki, Andrew Hill, Elena Engel, Ann Granchelli, Gabe Lockhart, Farnoush Banaei-Kashani, Russell Bowler

**Affiliations:** University of Colorado Denver; Colorado School of Public Health; National Jewish Health

## Abstract

1

**Background:** Chronic obstructive pulmonary disease (COPD) affects 5-10% of the adult US population and is a major cause of mortality. Acute exacerbations of COPD (AECOPDs) are a major driver of COPD morbidity and mortality, but there are no cost-effective methods to identify early AECOPDs when treatment is most likely to reduce the severity and duration of AECOPDs.

**Methods:** We conducted the first long-term (> 12 months), real-time monitoring studies of AECOPD with wearable sensors and self-reporting. We applied a deep learning-based autoencoder for feature extractions, then applied K-means clustering to detect heterogeneity. Accordingly, we proposed a weakly supervised active learning framework to develop anomaly detection models for robust identification of early AECOPD, and a clustered federated learning approach to personalize the anomaly detection models for early detection of heterogeneous subtypes of AECOPD. We evaluated this model by comparing it with other unsupervised learning models and federated learning models.

**Findings:** We identified two clusters based on the Silhouette score and SHAP analysis.One cluster shows high heart rate, low calories, and low steps; and the other has opposite characteristics. We also found out that a single subject could have exacerbation events from both clusters, indicating that there is not only subject-level heterogeneity but also event-level heterogeneity. Our weakly supervised framework outperformed unsupervised methods by 0.06 in average precision with 25 human annotation labels per subject. Our federated learning framework outperformed standard federated learning methods by 0.14 in F1 score and 0.17 in average precision.

**Interpretation:** We showed subject-level and event-level heterogeneity in AECOPD using mobile and wearable device data and developed a practical AECOPD detection framework with limited human annotated labels and keeping data private in each device.

## 2 Introduction

Chronic Obstructive Pulmonary Disease (COPD) is a leading cause of mortality in the United States and acute exacerbations of COPD (AECOPDs) are the major COPD morbidity that leads to excessive mortality, disease progression, and cost of COPD care. In a 5-year longitudinal follow-up study, we found each exacerbation was associated with a 30-75 ml/year decline in lung function, suggesting that AECOPDs contribute to COPD disease progression [1]. Exacerbations also decrease the quality of life [2; 3] and increase mortality [4; 5; 6]. Thus, there are still unmet needs in treating AECOPDs.

Treatment of AECOPDs is typically with corticosteroids and antibiotics. A small study showed that high intensity, supervised monitoring can lead to earlier treatment and better outcomes. Thus, the ability to predict and treat exacerbations early could likely modify the disease progression, reduce cost, and improve quality of life [7; 8]. Other studies have shown that the median time from AECOPD onset to treatment is 4 days and that early treatment is associated with faster recovery and less emergent hospitalization [9]. Unfortunately, 50% to 70% of AECOPDs are unreported [9; 2; 10; 11], suggesting that both late and unreported AECOPDs are unsolved clinical problems. Although it is generally accepted that early treatment of AECOPDs reduces the burden of hospitalization, little is known about the earliest, pre-health care utilization (pre-HCU) characteristics of AECOPDs, and there are no cost-effective methods for early at-home AECOPD detection in the week that precedes a HCU. This gap could be addressed with new wearable sensors and smartphone applications that can generate real-time data streams to monitor COPD patients at home; however, there is a need for automated tools that can process these data and develop personalized risk prediction models for early AECOPD identification. Furthermore, our preliminary work demonstrates that there is considerable heterogeneity in the clinical features that precede AECOPDs before healthcare utilization, which makes it even more difficult to identify AECOPD events before they happen.

Another barrier to improving treatment and outcomes of AECOPDs is difficulty in identifying an early AECOPD while the patient is still at home. The major drawback to home-based AECOPD disease management programs are that they require active participation and expensive personnel costs. An alternative solution to these programs could be automated data collection with wearable sensors and smartphone apps coupled with artificial intelligence/machine learning (AI/ML) based monitoring. However, the potential of clinical tools utilizing state-of-the-art AI/ML techniques has not yet been explored. In this study, We aimed to analyze the heterogeneity of AECOPDs and evaluate a deep learning framework for anomaly detection with minimum labeled data and privacy data protection.

The goal of this project is to identify the heterogeneity of COPD in data coming from wearable devices and introduce methods for advanced and robust detection of early clinical features in heterogeneous subtypes of AECOPD. If successful, our proposed methods can also be adopted for the early detection of acute exacerbations in other chronic diseases.

## 3 Methods

### 3.1 Recruitment of subjects

Subjects were recruited from a single special clinic and through online advertising. Eligibility included a physician diagnosis of COPD and an AECOPD treated with corticosteroids or antibiotics in the preceding 12 months in order to enrich for future exacerbations. This study was approved by an institutional review board and all subjects gave informed written consent.

### 3.2 Study procedures

193 subjects were screened and 73 subjects consented. After completing informed consent, 50 subjects were enrolled in a minimum 1 week run in period in which they were asked to complete the The Exacerbations of Chronic Pulmonary Disease Tool (EXACT) on their smartphone (Apple or Android). The EXACT is a 14-question patient-reported outcome (PRO) measure for standardizing the symptomatic evaluation of AECOPDs and acute exacerbations of chronic bronchitis (AECB) in natural history studies and clinical trials [12]. Fifty subjects completed the EXACT for at least 6 days in the first week of the run-in and were mailed the following sensors: a smart watch (Fitbit Sense 2 [13] or Apple Watch series 6 [14]), inhaler sensors (Propeller Health [15]), and a smart thermometer (Kinsa Health [16]). Subjects were then guided through the installation of the devices and applications over the phone and asked to wear the smartwatch and answer the smartphone survey daily.

### 3.3 Event adjudication

Subjects were encouraged to self-report AECOPDs, but were also called every 4 months by a research coordinator to complete an exacerbation questionnaire. If necessary, AECOPD medical records were obtained, redacted, and reviewed by a pulmonologist, who adjudicated (1) whether the event was an AECOPD and what date it started; and (2) the severity of the event: severe being defined as requiring hospitalization or emergency room; moderate being not severe, but requiring HCU with corticosteroids or antibiotics; or mild, not requiring HCU or prescribed medications.

### 3.4 Data Collection and Processing

Subjects were asked to complete the digital EXACT at a consistent self-selected time during the evening. For subjects with an iPhone, data were collected using a bespoke iOS app [17]. For other subjects, we collected smartwatch data using Fitabase [18] and the EXACT using RedCAp [19]. Propeller data were collected from the Propeller research portal. Repeat actuation occurring < 2 minutes apart were considered as a single inhaler use event [20]. Physiologic data including heart rate (HR), steps, EXACT, inhaler usage, and adjudicated AECOPD events were then temporally aligned.

### 3.5 Predictor Variables

The predictors/features we used are summarized in Table 1 and 2. We converted all features into a daily basis by averaging for the purpose of computational efficiency.

**Table 1.** Description of data features.

| Feature | Description |
| --- | --- |
| EXACT | Development of the EXAcerbations of Chronic Obstructive Pulmonary Disease Tool |
| Oxygen Saturation | A measure of how much hemoglobin is currently bound to oxygen compared to how much hemoglobin remains unbound. |
| Rescue Inhaler Use | Medication for relieving symptoms |
| Controller Inhaler Use | Medication for prevention |
| Heart Rate | The number of times your heart beats per minute |
| Steps | How many steps you take each minute |
| Calories | Total calories burned for the min [12] |

**Table 2.** Details of data features.

| Feature | Domain | Source | Type of Data(Range) | Temporal Resolution |
| --- | --- | --- | --- | --- |
| EXACT | Symptoms | App Survey | Score(0-100) | Daily |
| Oxygen Saturation | Physiology | App Survey | SpO2(0-100%) | Daily |
| Rescue Inhaler Use | Medication | Propeller | Puffs per day | Daily |
| Controller Inhaler Use | Medication | Propeller | Puffs per day | Daily |
| Heart Rate | Physiology | Fitbit or Apple watch | Average beats/min | min |
| Steps | Activity | Fitbit or Apple watch | steps | min |
| Calories | Activity | Fitbit or Apple watch | calories | min |

### 3.6 Data analysis

Our analysis process included data preprocessing, dimensionality reduction, clustering, and SHAP values analysis [21] as described below.

#### Missingness and training dataset

We selected 8 patients who had at least 2 exacerbation events, in order to make both train datasets and test datasets for anomaly detection experiments. Thereafter, we converted all features into daily basis data by averaging for computational efficiency. We dropped consecutive missing values with > 1 consecutive day of missing data for heart rate and steps while we dropped missing values with > 3 days for EXACT and oxygen saturation, and we imputed missing values with linear interpolation otherwise. We standardized each feature after imputation.

#### Dimensionality reduction

Because daily-based data is still high dimensional and it is hard to compute the distance between those data, we reduced data dimensionality by using an LSTM-based autoencoder. We converted all features into 8-dimensional vectors. (Figure 2) When we focused on patient-level heterogeneity in this section, we selected one consecutive pre-exacerbation event data group from each patient to observe data trends related to exacerbation events. *Clustering*. We applied K-Means clustering [22] to the LSTM [23] embedded data and analyzed silhouette score [24] and analyzed each cluster by game theory-based Shapley Values (SHAP Values) [21]. We selected the number of clusters *k* = 2 because 2 clusters led to the highest silhouette score (6).

#### Early AECOPD prediction

We trained an LSTM-based classification model to test whether it could detect anomalies (AECOPD) from in the 0-30 days prior to HCU from an AECOPD. We designed an anomaly detection framework for the purpose of early detection of AECOPD. The overview of our framework is summarized in Figure 3. This framework consists of two components. The first component is an active learning-based anomaly detection component in each patient. This allows each subject to train an anomaly detection model within their device without any privacy concerns or communication costs with a central server. The second component is Federated Learning (FL) [25]. FL allows each patient to communicate with a central server and share knowledge with other patients without sharing raw data, thus keeping individual data private. This is possible by sharing only trained model parameters. Because we had observed AECOPD heterogeneity by subject, we considered heterogeneity during the server aggregation. We applied the same data preprocessing process as figure 1. First, we randomly selected one exacerbation event per patient for test data, and then we assigned 80 % of the data as the training dataset and 20 % as a test dataset. We used a modified version of Label-Efficient Interactive Time-Series Anomaly Detection (LEIAD) [26]. This framework utilizes active learning to maximize the model performance with a limited number of human-annotated label data with an ensemble of unsupervised anomaly detection, which increases the robustness of the active learning label generation process. Our model considers multi-variate time series data and uses an LSTM-based deep learning model to be able to capture complex multi-variate data structures. In our experiment, we added 5 true labels after each iteration.

**Fig. 1.**
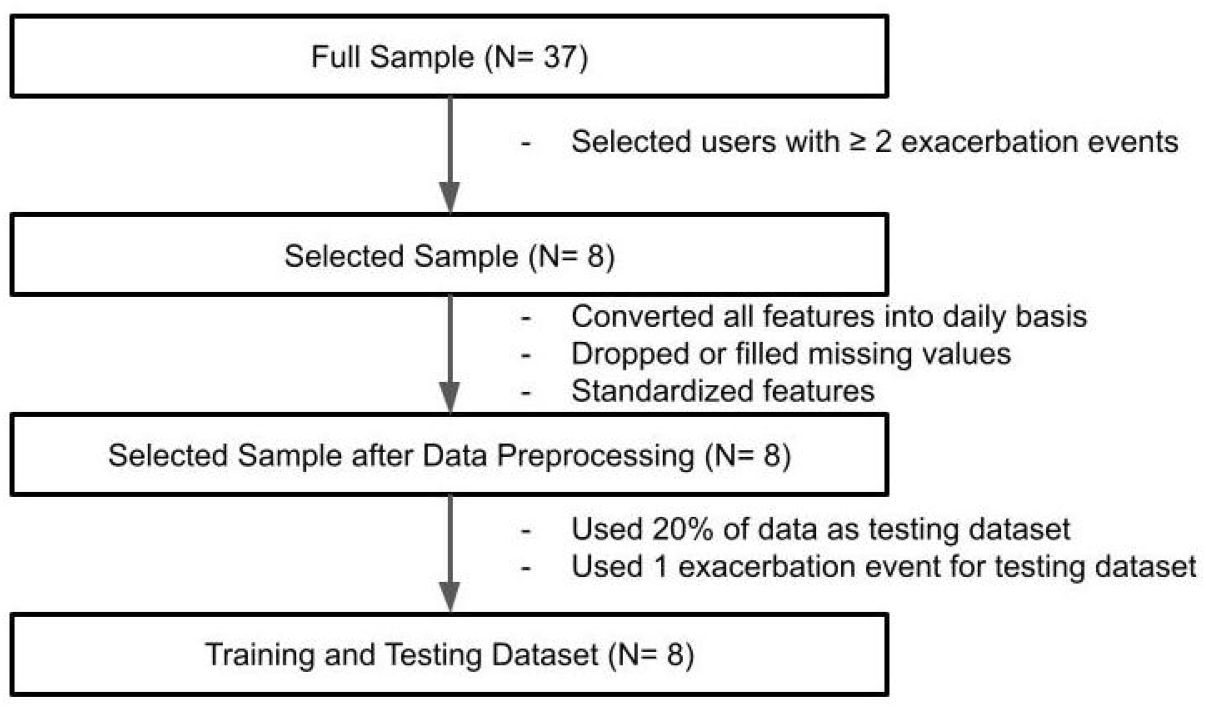
Data preprocessing flow.

**Fig. 2.**
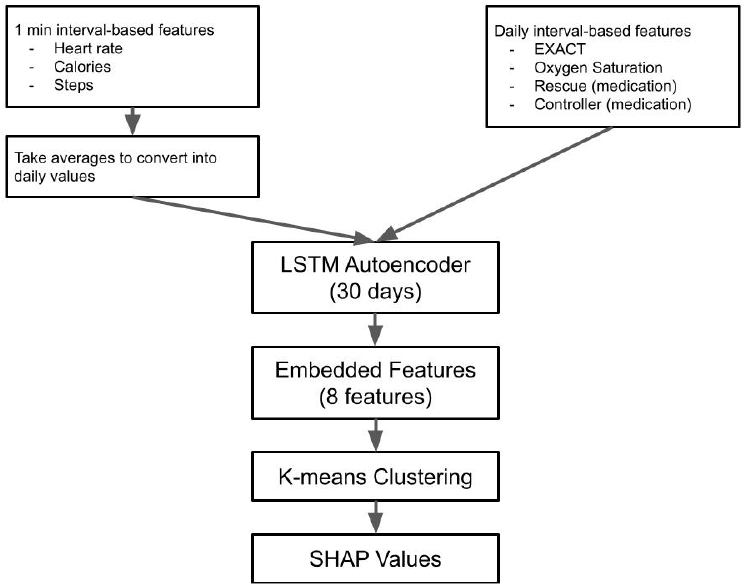
Dimensionality reduction flow.

**Fig. 3.**
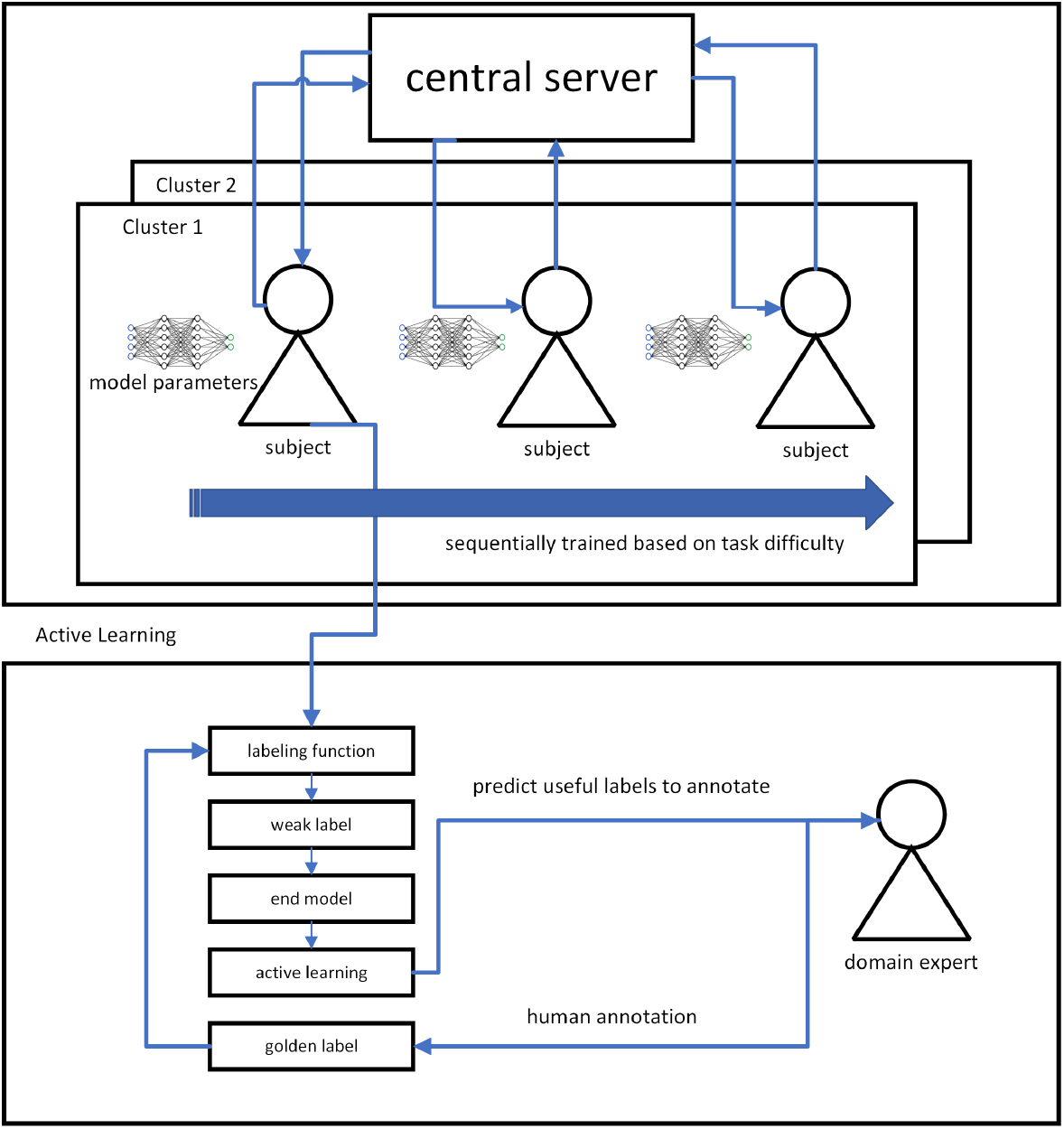
Framework Overview.

#### Federated Learning (FL)

FL is a relatively new machine learning paradigm where multiple machine learning models train collaboratively without sharing raw data. This is possible by sharing only parameters with a central server, and the server aggregates those parameters by a computational function, such as averaging or weight averaging, to share knowledge implicitly. One of the main challenges in applying FL to our setting is data heterogeneity. As we have seen, each patient, and even each exacerbation event, could be heterogeneous. Hence, aggregating all patients’ models into a single global model could be detrimental. The other potential issue is paucity of data. Each patient has only 2-4 exacerbation events, which could lead to poor performance in each model, and aggregating poor local models does not usually work.

To overcome these challenges, we proposed a novel deep learning-based method with three components. The first one is the daisy-chain algorithm [27] which allows a central server to send a global model to one subject, train the model, send it back to the server, and iterate this process for another subject. This replicates training using all raw data, which works well when data are limited. The second component is clustering based on a model loss, which is different from a common distance measure such as cosine similarity of model parameters. In our case, since the data from each patient device is limited, each local model might be poor, which might not represent data characteristics well. Therefore, we used model loss as a distance measure. First, we randomly select one subject and train a local model. Thereafter, we send the model to all other patients and compute model loss, and if the model loss is smaller than a threshold, we make a cluster for them. If a model loss is smaller, that means that a model that works for a local dataset could work for another dataset, implying similar data distribution. In addition to that, this can measure data characteristics more directly than using cosine similarity for model parameters. That is why our similarity measure is more suitable for limited datasets than cosine similarity would be. The third component is curriculum learning [28]. Curriculum learning is a machine learning technique where we first train a model using easier data/tasks and later train a model with more difficult data/tasks, which is similar to how human brains work. In order to optimize the order in the daisy-chain algorithm, we used curriculum learning.

### 4 Results

We summarized the consort diagram in the Figure 4, Characteristics of subjects in Table 3, and data statistics in the Table 4. 37 of 193 subjects were used for data analysis. There were 20 moderate and 5 severe exacerbations. Exacerbation rates were 0.58 per year.

**Table 3.**
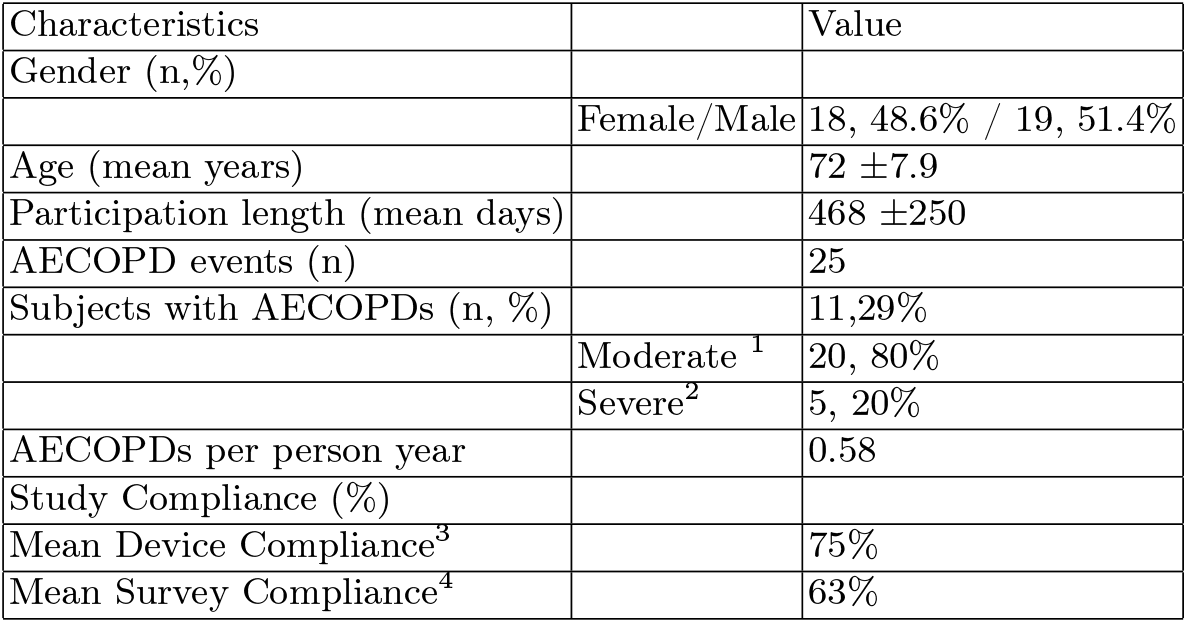
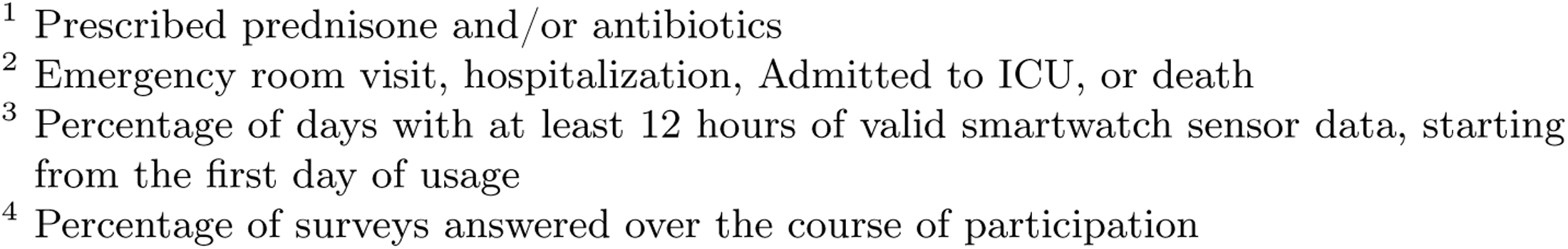
Characteristics of subjects.

| Characteristics |  | Value |
| --- | --- | --- |
| Gender (n,%) |  |  |
|  | Female/Male | 18, 48.6% / 19, 51.4% |
| Age (mean years) | | 72 ±7.9 |
| Participation length (mean days) | | 468 ±250 |
| AECOPD events (n) |  | 25 |
| Subjects with AECOPDs (n, %) |  | 11, 29% |
|  | Moderate <sup>1</sup> | 20, 80% |
|  | Severe<sup>2</sup> | 5, 20% |
| AECOPDs per person year |  | 0.58 |
| Study Compliance (%) |  |  |
| Mean Device Compliance<sup>3</sup> |  | 75% |
| Mean Survey Compliance<sup>4</sup> |  | 63% |

**Table 4.** Data statistics.

| Feature | mean ± standard deviation | Missing rate |
| --- | --- | --- |
| EXACT | 38.5±12.7 | 23.7 |
| Oxygen Saturation | 93.0±2.80 | 52.4 |
| Rescue Inhaler Use | 1.56±3.40 | 0 |
| Controller Inhaler Use | 1.67±2.73 | 0 |
| Heart Rate | 76.4 ±13.3 | 54.7 |
| Steps | 1.99±8.62 | 46.4 |
| Calories | 1.20±0.85 | 46.4 |

**Fig. 4.**
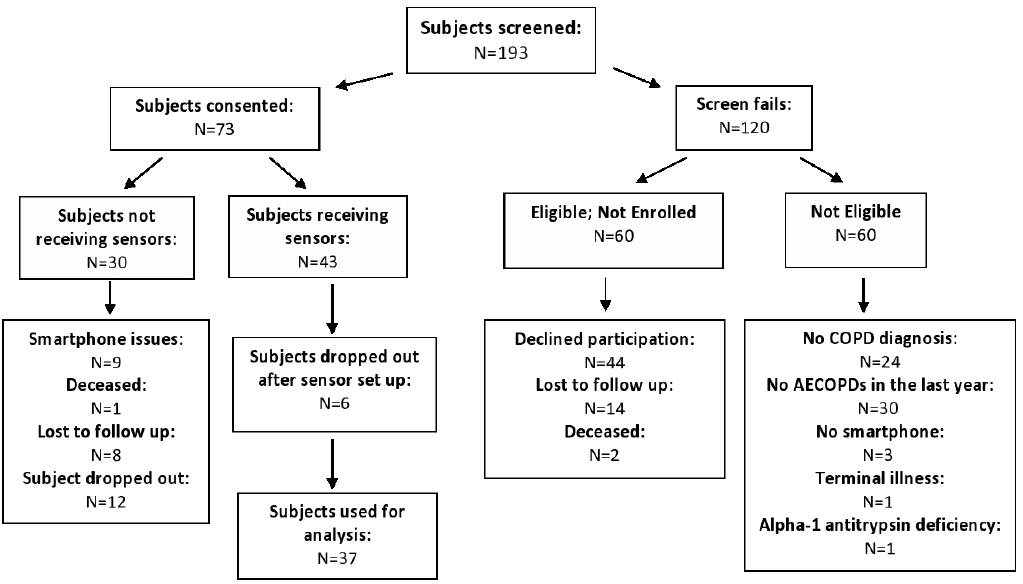
Consort Diagram.

We selected the number of clusters *k* = 2 for patient-level clustering because it achieved the highest silhouette score and steep slope in the Elbow method (Figure 5 and Figure 6). The silhouette score near 0.5 shows that the 2 clusters are somewhat separate, which may indicate that there is some heterogeneity among data trends before exacerbation events among patients.

**Fig. 5.**
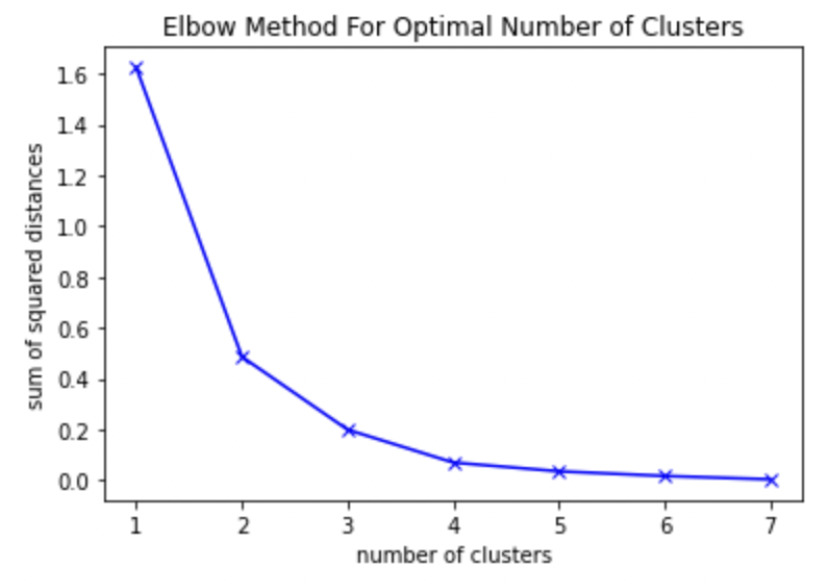
Elbow method for patient-level clustering.

**Fig. 6.**
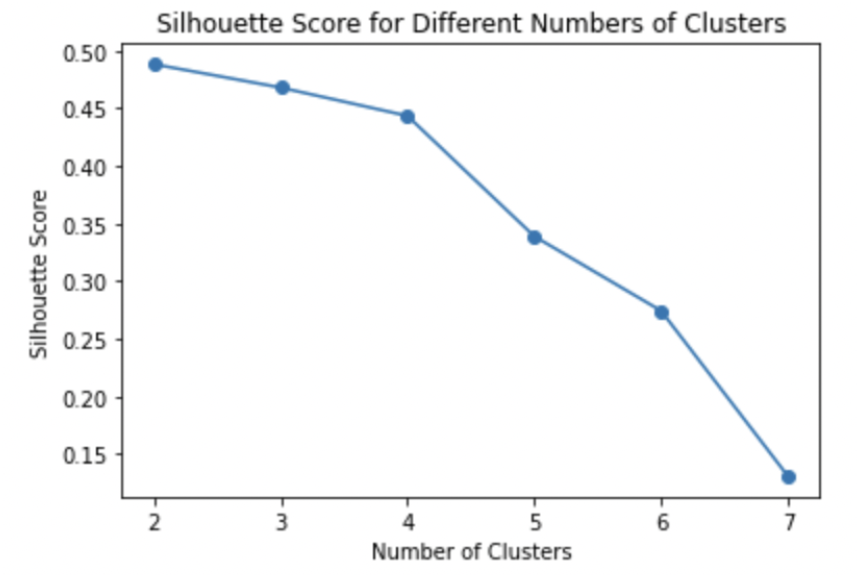
Silouette score for patient-level clustering.

In addition to the silhouette score, we analyzed each cluster by game theory-based Shapley Values (SHAP Values) to observe the characteristics of each cluster. High SHAP values of features mean that those features are more informative as predictive variables. In our subject-level analysis, we visualized SHAP values of clustering. Hence, each SHAP value shows the contributions of each feature to clustering. Figure 7 shows absolute values of SHAP values for 30 days before an exacerbation event. Figure 7 implies that predictive variable data has heterogeneity from day 16 to day 30 where day 31 is the day when an exacerbation event occurs. SHAP values increase over time, meaning that data closer to an exacerbation event has more impact on clustering. This means that data trends could show more unique trends in each cluster as it gets closer to an exacerbation event. This shows the heterogeneity of exacerbation events and the necessity of anomaly detection considering the heterogeneity among patients.

**Fig. 7.**
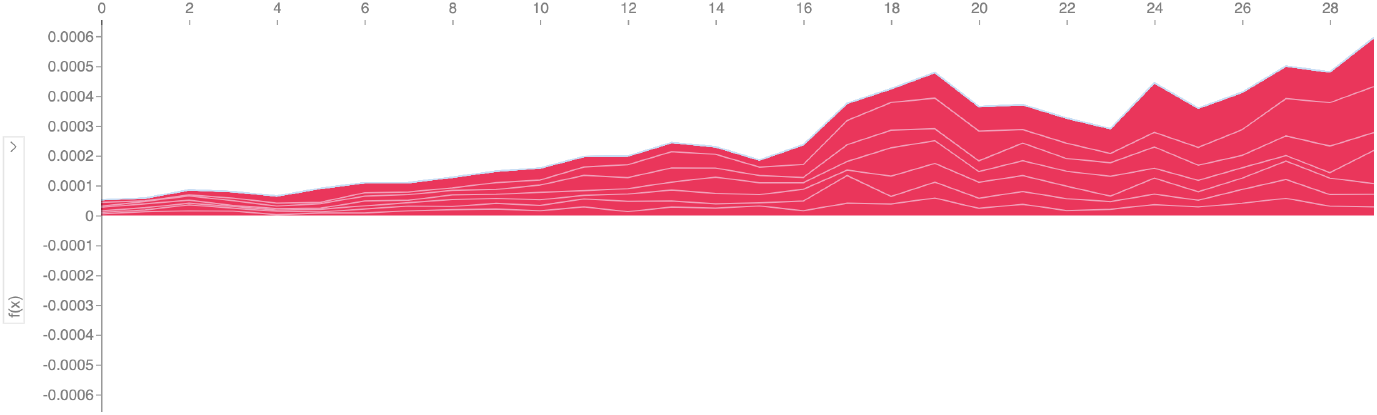
Shap values for 30 days before an exacerbation event (subject-level)

Figure 8 shows how each feature contributes to clustering. Figure 8 shows that one cluster (namely, cluster 0) has low steps, low calories, high EXACT, high heart rate, high control medication, high oxygen saturation and the other cluster (namely cluster 1) has low exact, high oxygen, and low rescue. These results with the silhouette score analysis indicate that data trend before an exacerbation event has heterogeneity and the heterogeneity increases over time as it gets closer to an exacerbation event.

**Fig. 8.**
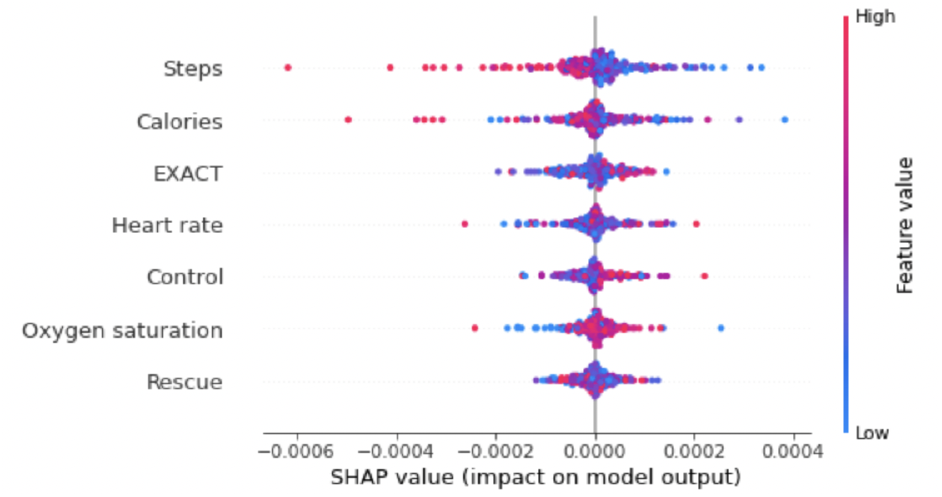
Shap values (subject-level)

Our SHAP analysis for clustering exacerbation events regardless of subjects (event-level) is in figure 9 and 10. Figure 9 shows a similar trend as subject-level heterogeneity analysis, which shows an increase in heterogeneity over time. Figure 10 shows that one cluster (cluster 0) has high heart rate, low control medication, low steps, low EXACT, low rescue medication, low calories and high oxygen saturation, and the other cluster (cluster 1) has opposite characteristics. When we compare the top 3 impactful features of subject-level and event-level clusters (3 features from the above in figure 8 and 10), cluster 0 in both clusters shows high heart rate, low calories, and low steps while EXACT and control medications do not match. This result shows we could observe a similar cluster of high heart rate, low calories, and low steps in both subject-level and event-level clustering.

**Fig. 9.**
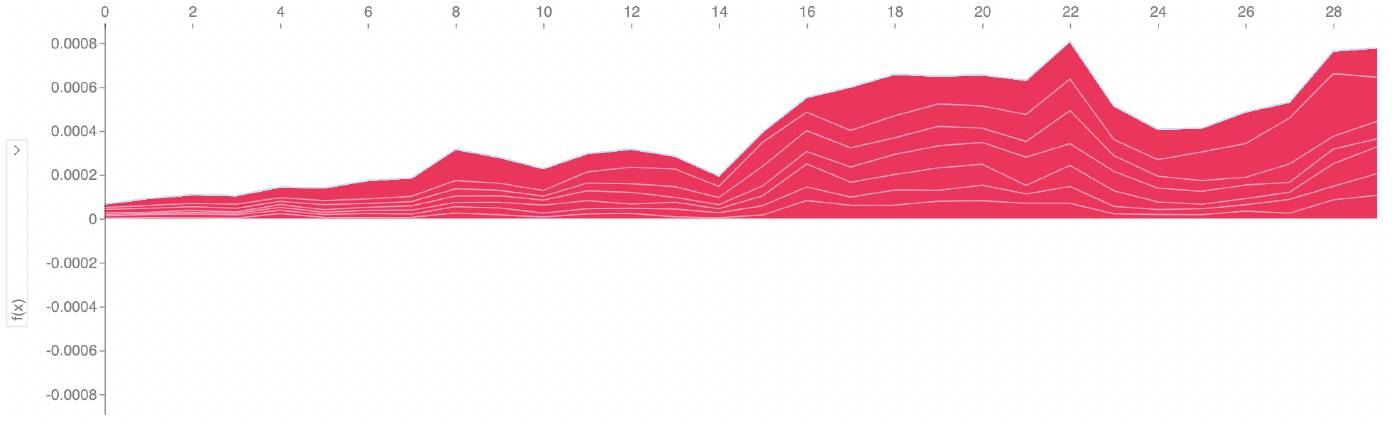
Shap values for 30 days before an exacerbation event (event-level)

**Fig. 10.**
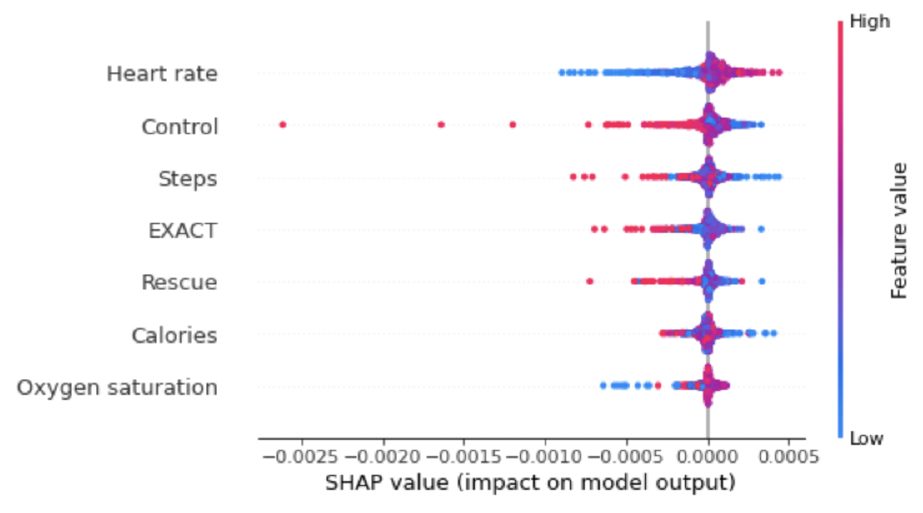
Shap values (event-level)

Table 5 shows which cluster each exacerbation event belongs to. 5 out of 8 patients have 2 clusters in a single patient, indicating that there could be heterogeneity even among a patient.

**Table 5.**
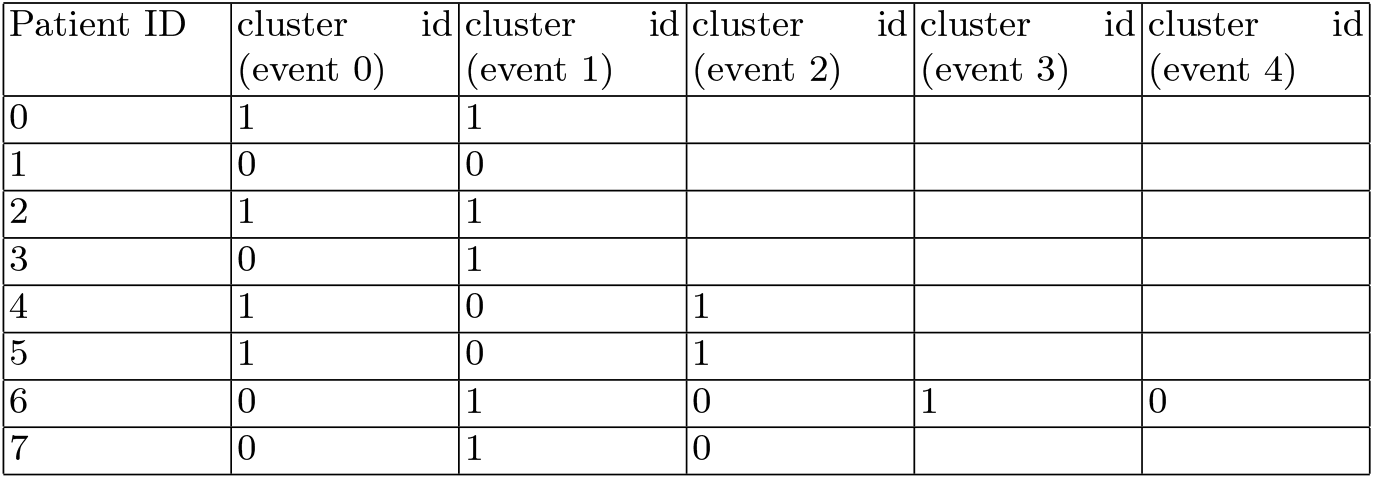
Characteristics of study participants.

Figure 11 and figure 12 are data samples. In figure 11, we can observe similar data trends before exacerbation events (yellow areas) where EXACT, rescue medications, heart rate, and calories increase. This implies that this subject does not have heterogeneity in AECOPD. In figure 12, we can observe high oxygen saturation, high calories before one exacerbation event whereas we can observe low oxygen saturation, low heart rate, low steps and low calories before the other exacerbation event, indicating there is heterogeneity in AECOPD in a single subject. This example also shows the existence of event-level heterogeneity.

Figure 13 and 14 show performance comparison in F1 and Average precision regarding LEIAD and other unsupervised learning models in the test dataset. They show average results among 8 patients. We observed that our modified LEIAD achieved the highest performance in Average Precision after 3-5 iterations. This indicates that LEIAD could outperform unsupervised models only with 15-25 human-annotated labels per patient. This was possible because LEIAD identifies which label data we need to improve model performance with a limited number of labeled data. Our modified LEIAD achieved the second-best result in F1 score and we noticed the performance drop after 25 human-annotated labels. This performance drop could be explained by the fact that there was heterogeneity in labels because our algorithm added those true heterogeneous labels.

**Fig. 11.**
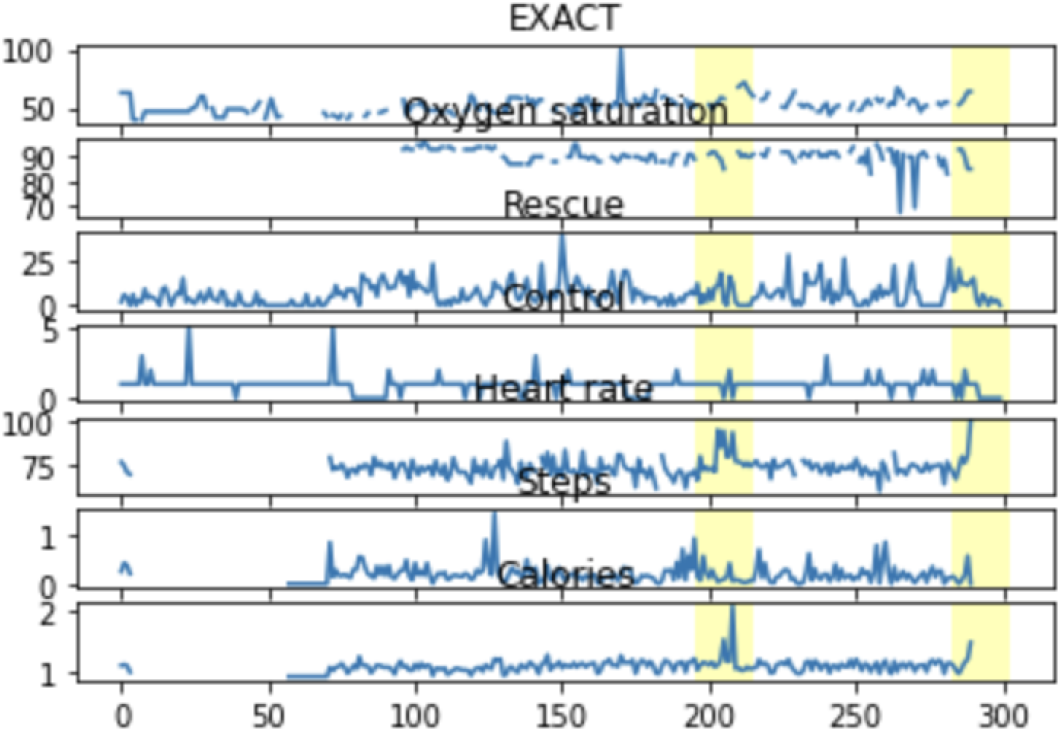
Sample data 1 (no heterogeneity in AECOPD)

**Fig. 12.**
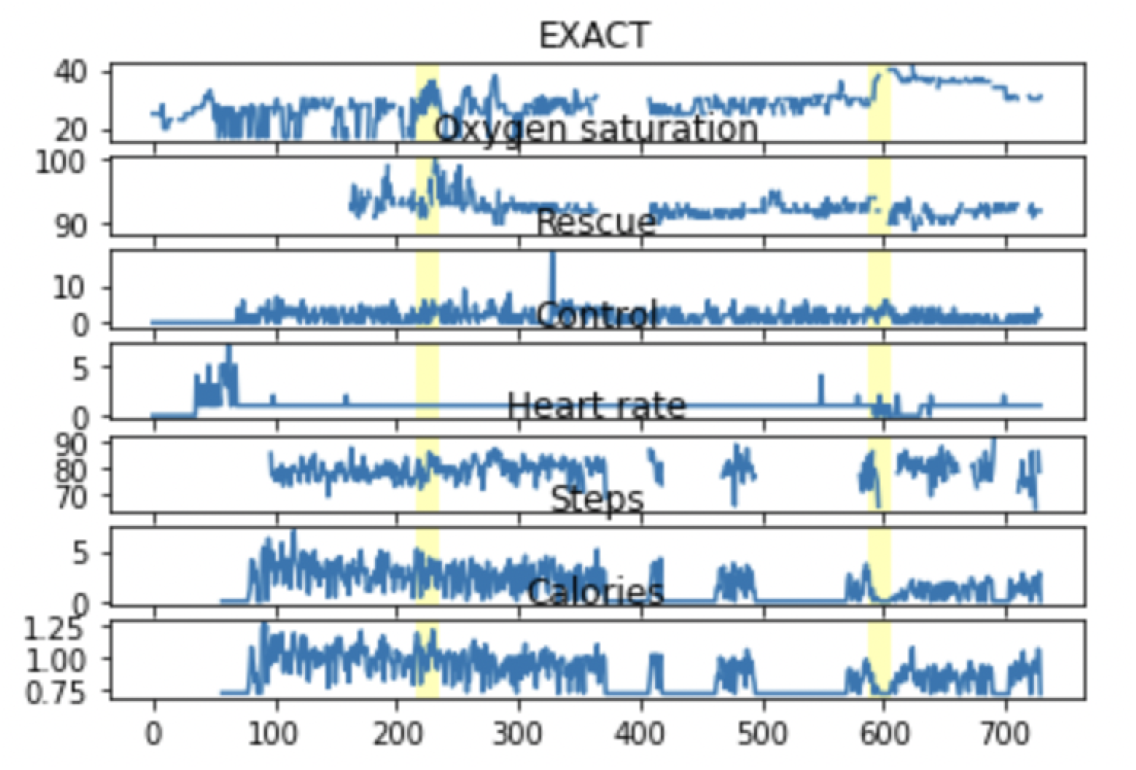
Sample data 2 (heterogeneity in AECOPD)

**Fig. 13.**
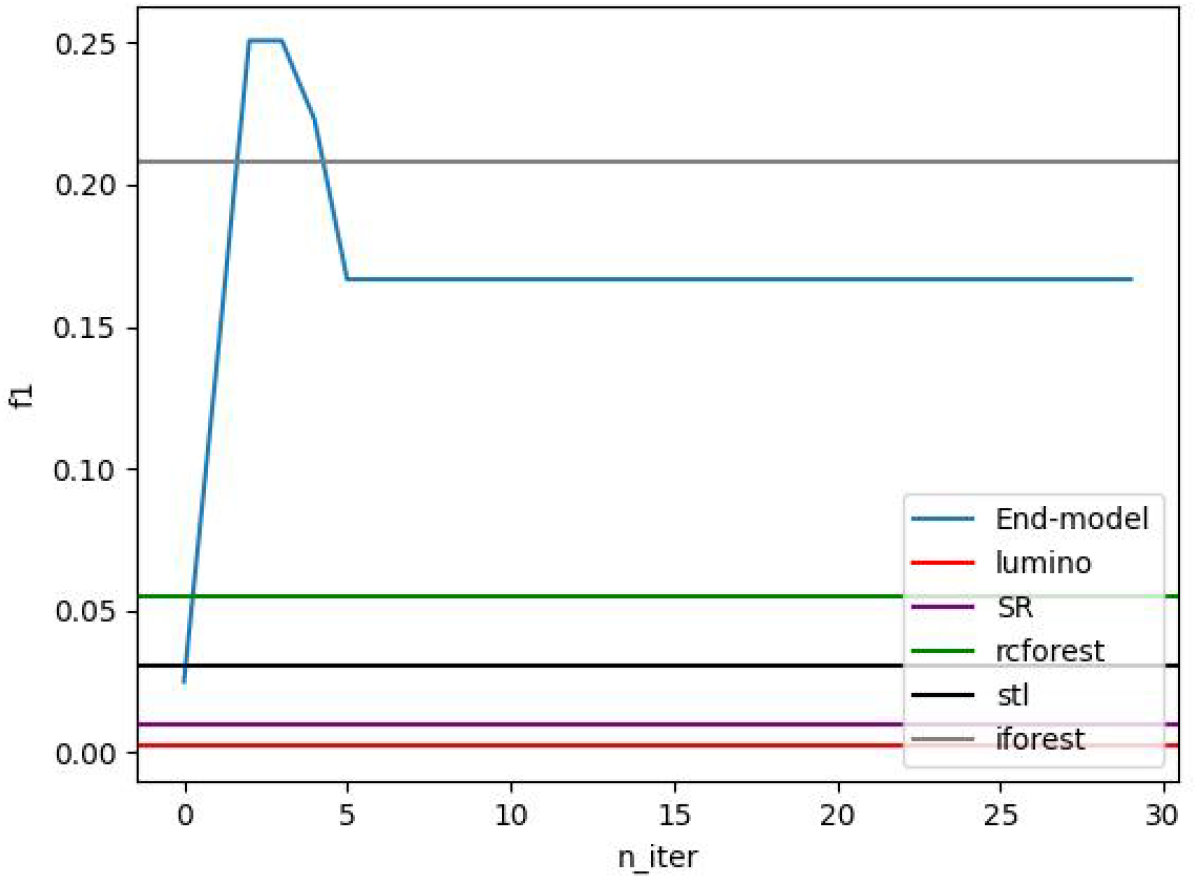
Average F1 score among patients of LEIAD and other unsupervised methods.

**Fig. 14.**
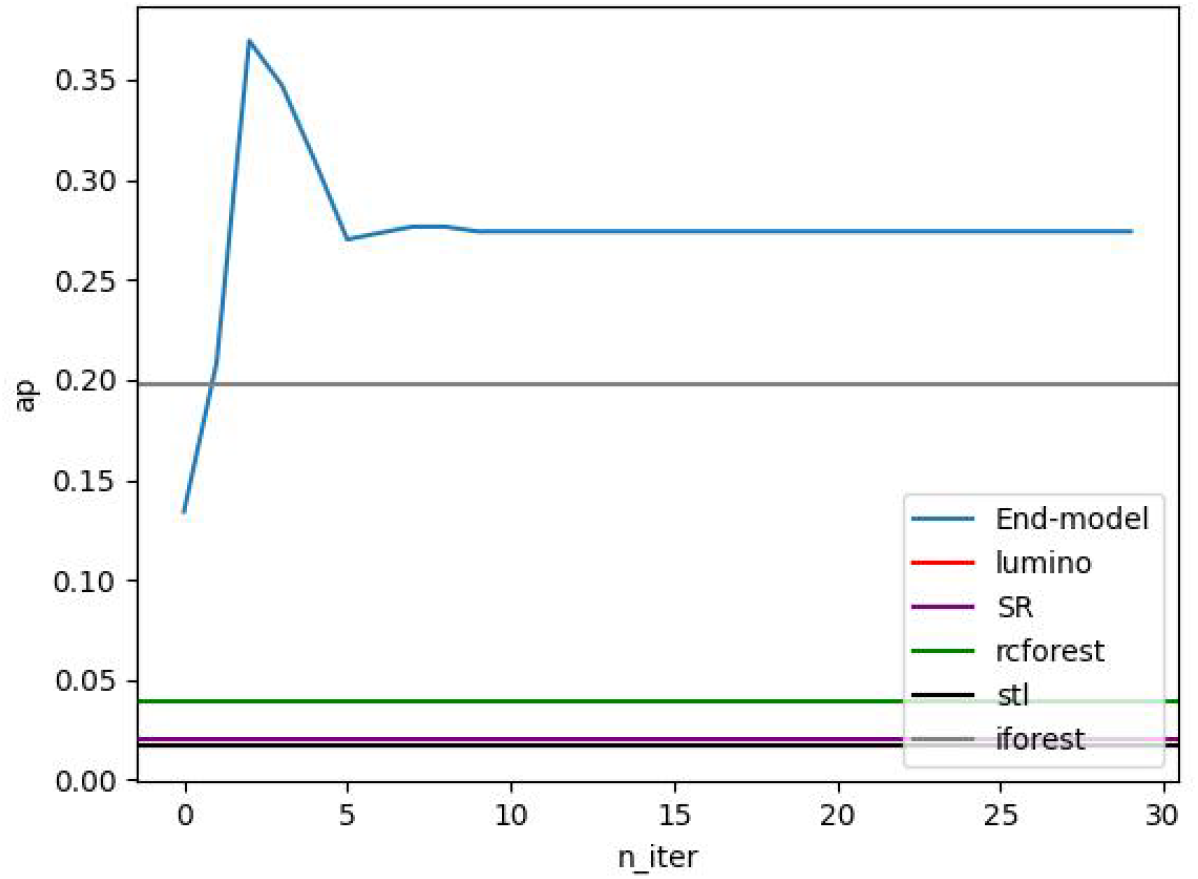
Average Average precision among patients of LEIAD and other unsupervised methods.

Table 6 shows the results of various FL algorithms for our dataset. We first used LEAID with 25 human annotations and then applied each FL algorithm to test practical applications. LEIAD + Standalone, LEIAD + FedAvg, and LEIAD + FedDC achieved the same results, which indicates that FL does not improve the results. This could be explained by that aggregating all local models is not beneficial due to data heterogeneity. When we add clustering and curriculum learning, those models increase model performance, indicating that aggregating models among clusters and curriculum learning would be effective when there is heterogeneity and data is limited. This is because sharing common knowledge only within the same clusters is beneficial in patients in the same clusters. The biggest performance gain was possible by clustering, also showing heterogeneity in AECOPD.

**Table 6.**
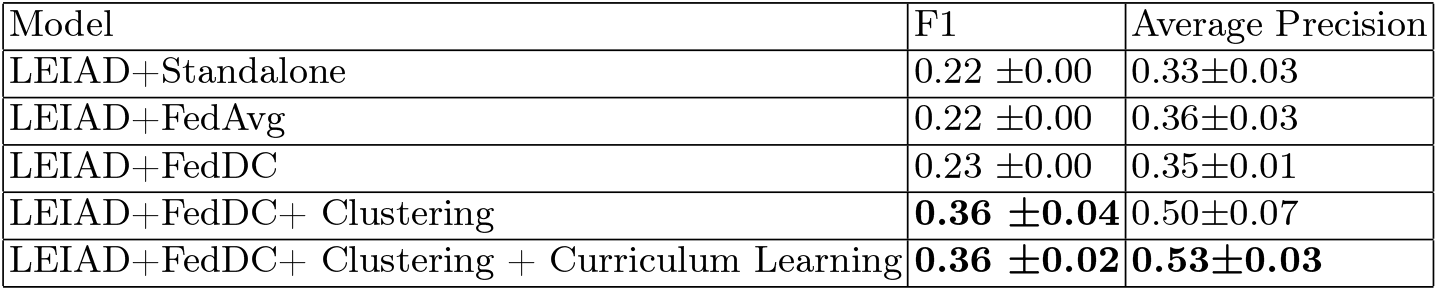
Model performance comparison.

## 5 Discussion

In this work, we analyzed heterogeneity in AECOPD at the subject-level and event-level, and we proposed a practical anomaly detection framework based on weakly supervised learning and federated learning. We found that the data trend right before an exacerbation event has 2 clusters, which indicates that there is AECOPD heterogeneity. We also observed that some subjects have exacerbation events from both clusters, meaning that there is heterogeneity even among a single subject. In order to address this heterogeneity and lack of data, we proposed an anomaly detection framework utilizing weakly supervised learning and federated learning. Our weakly supervised learning method helps medical experts to make annotated data efficiently so that we are able to maximize model performance with limited human annotated labels. Our Federated Learning method allows us to make patient-level clusters and share knowledge within clusters without sharing raw data (for privacy protection). Our experiments show the potential of the practical application of the framework in a real-world setting where medical experts add a small number of annotated labels, train models in each patient device, and apply federated learning in a central server for further improvement considering heterogeneity.

We explored heterogeneity in AECOPD and found important insights about patient-level and event-level heterogeneity. We found two clusters. One cluster shows high heart rate, low calories, and low steps, and the other has opposite characteristics. Interestingly enough, a single subject could have exacerbation events from both clusters, meaning that this AECOPD heterogeneity is not necessarily subject-specific, which makes early detection more challenging. SHAP values also showed that the heterogeneity increased over time, starting to show a signal 10-20 days before an exacerbation event, and becoming stronger over time. This may show the possibility of earlier detection. However, more exacerbation training data sets will be needed for earlier detection.

We proposed a practical anomaly detection framework and showed potential through several experiments. Each subject utilizes active learning where an ensemble of unsupervised models suggests the most informative data to annotate. We found that our model outperformed baseline solutions based on unsupervised anomaly detection only with 15-25 annotated events, which shows practical use with high efficacy. We modified the existing framework to adjust to our multivariate and limited data scenarios by including an over-sampling process, deep learning-based feature extraction, and an ensemble of all features. We also used federated learning in order to share knowledge among patients without sharing raw data. FL achieves this goal by only sharing model parameters. In addition to that, we applied clustering based on model loss where we trained one model with one patient data and computed model loss for each patient and group patients with small loss. Thereafter, we used a daisy-chain algorithm and curriculum learning to address the problem of a limited number of events. Training a model sequentially, especially in order of task difficulty, replicates a situation where we train a single model with all local data, which leads to model performance improvement in a limited number of data.

In addition to early detection, efficient home monitoring could ameliorate disparities in health care. There are also health disparities in COPD (and in turn, in AECOPD). The three leading causes of death in rural areas (heart disease, cancer, and stroke) have been decreasing, yet the rural pre-COVID mortality rate for COPD has been increasing [29]. Rural patients have significantly higher COPD prevalence, Medicare hospitalization for COPD, and COPD related deaths [30; 31; 32]. Black Americans are also a group for which there is poor access to COPD special care and worse quality of life with exacerbations [33]and also reduced access to special care for chronic lung diseases compared to white Americans [34].

There are some limitations in this work. First, we converted minute-based data coming from Fitbit and Apple Watch, such as heart rate and steps, however, this could have led to some information loss, leading to model performance degradation. On the other hand, training a model using minute-based data requires much more computational power, so our future work is to develop a computationally efficient method so that we can realize it in each patient’s device. Second, the variety of data features is still limited. For instance, we are considering collecting temperature and air quality data using GPS. Third, since most of the patients have only 2 exacerbation events, they have only 1 exacerbation event for training data, hence, we were not able to develop a framework and test event-level heterogeneity. Our current framework only considers patient-level heterogeneity, therefore, our future work is to collect more data per patient and extend our method to handle event-level heterogeneity.

Our work detected heterogeneity at the subject-level and event-level and proposed a practical AECOPD detection framework with limited human annotations and privacy protection. We identified two distinct clusters and found that a single patient could have an exacerbation event from both clusters. Our AECOPD detection framework incorporates weakly supervised learning and federated learning with novelty in accommodating multi-variate time series data, clustering based on local data implicitly, and applying FL in the order of task difficulty. We showed improvement from our baseline solutions and showed potential for practical medical applications. Our research shed light on new research directions in AECOPD heterogeneity and how to address it in real-world applications with recent artificial intelligence technologies.

