## Supplemental Document for "Heterogeneity analysis of acute exacerbations of chronic obstructive pulmonary disease and a deep learning framework with weak supervision and privacy protection"

Table of Content

1 Data Collection
2 Imputation
3 Early detection
4 Anomaly Detection Framework

### 1 Data Collection

Respiratory symptoms such as dyspnea, wheezing and sputum production define AECOPDs, but are traditionally ascertained retrospectively. In our pilot studies we used diaries such as the EXAcerbations of Chronic Pulmonary Disease Tool (EXACT), which more reliably assesses frequency, severity, and duration of COPD exacerbations [1] and can be administered electronically using smartphones [2; 3]. Physical activity and rescue medication use are also two important factors that are strong predictors of COPD outcomes but are less reliable when self-reported. Recent technological advances now allow real time passive electronic collection of activity, heart rate, pulse oximetry, temperature, and inhaler use (see Figure 1). These sensors are particularly attractive to use because they passively collect and transmit data through smartphone. Using aforementioned data collection means, we have conducted two pilot studies of 3-week and 12+ month enrollment duration to collect data and evaluate efficacy of basic methods as well as challenges in early detection of AECOPD. First, in our recently published 3-week pilot study , 184 COPDGene subjects were enrolled over six months at one clinical center. The subjects used an Android smartphone to complete an eDiary (EXACT), continuously wore an activity monitor (ActiGraph [4]) for monitoring steps and calories, and used real-time rescue inhaler monitoring (Adherium SmartInhaler) as well. Although the three-week pilot was not powered to detect health care utilization exacerbations, we detected 9 EXACT defined exacerbation events. We also identified novel rescue inhaler patterns that were independent of total inhaler doses per day and found that the EXACT score was positively associated with COPD progression and daily rescue inhaler use [2]. These publications demonstrated that some exacerbation events have prodromal symptoms that can be detected with real-time monitors while the subject is at home, but also suggest the need for the development of machine learning algorithms for better real time detection of events with integrated data. Full compliance (all three devices) over three weeks was 98% (180/184). Post study interviews with the subjects indicated that they did not find the study burdensome because most of the data are collected passively, other than the eDiary (which is short, and our subjects do not report as excessively burdensome. However, academic reviewers were skeptical that subjects would continue to wear devices and answer eDiary over 12 months. Therefore, for a second pilot study we enrolled new subjects from the community for a 12-month study. To date, this pilot study has enrolled 30 subjects at a single site (National Jewish Health) and only one subject has withdrawn from the study. Nearly all subjects have chosen to stay in the study for more than 12 months. In the 12-month pilot we used Fitbits instead of Actigraphs and we used Propeller instead of Adherium inhaler sensors. The COVID pandemic spurred additional innovations in the 12-month pilot including: electronic consent, remote enrollment, and addition of temperature and oxygen saturation to the sensor streams. In 2021 we added COVID surveillance in our three month follow up; however, none of the AECOPDs were due to COVID, which was consistent with marked reduction of AECOPDs in general because COPD patients were self-isolating in 2020 and 2021. In January 2022 Apple began providing us with the latest Apple watches (which can collect oxygen saturation data) and in April 2022 Apple approved our iPhone app for real-time collection and streaming of eDiary, inhaler usage and sensor data (steps, heart rate, oxygen saturation, and temperature) through Apply HealthKit to a protected cloud bucket (see Figure 2). Compliance with both eDiary and devices has been excellent in the 12-month pilot. For instance, 83% of subjects answered the EXACT survey on more than 75% of days and 74% of subjects wore the wrist device (Fitbit) more than 85% of days.

To begin to address this problem, we have conducted preliminary home-based real-time monitoring studies of high-risk COPD patients using wearable sensors. In our pilot work, we demonstrate a real-time, home-based monitoring approach using eDiaries such as the EXAcerbation of Chronic Pulmonary Disease Tool (EXACT) [5], real-time monitoring of inhaler and wrist sensors [2] and residential bioaerosol sampling, is feasible (>80% compliance) and can be cost-effective over both short term (3-weeks) and long-term (>12 months). These preliminary studies are the first to include and integrate real-time reporting of activity, heart rate, SpO2, temperature, inhaler use, and symptoms, and they have allowed us to develop and test basic unsupervised methods for early detection of AECOPD.

### 2 Imputation

One issue with the collected sensor data was considerable missing data. Participants may occasionally forget to utilize one of the devices or answer the survey, resulting in missing values for one or more sensor streams.

#### 2.1 Missing Values

Table 1 shows the pattern of missingness in the dataset. We observe heterogeneity of patterns of missingness for features, patients, and timing. Table 2 shows the correlation of missingness between every 2 features. It shows high correlation between EXACT and oxygen saturation, rescue and control medications, and heart rate and steps. All these pairs are collected from the same methods/devices, therefore, we could see that the way of collecting data has a big impact on missingness.

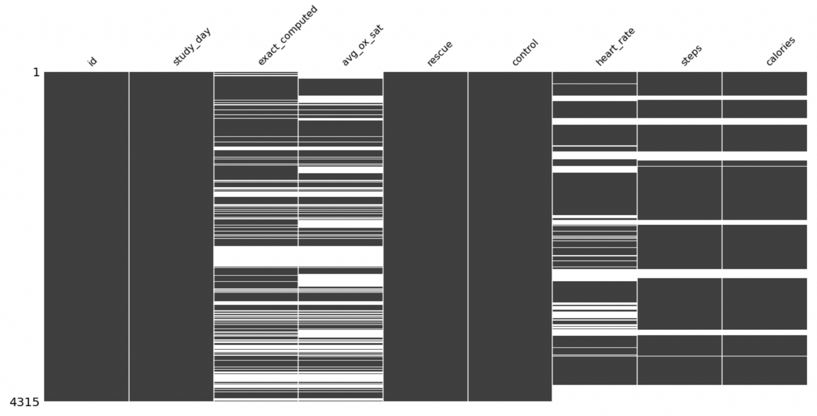


**Fig. 1.** Missing data distributions

#### 2.2 Dropping Missing Values

We used 2 strategies to handle missing values. The first strategy is to drop missing values based on a threshold. We need to find a threshold which does not drop too many missing values, but also removes data with too many missing values. In order to find an appropriate threshold to meet this requirement, we visualized the number of missing values for each consecutive day in figure 3 and 4 for oxygen saturation feature. In this case, we selected 4 days as a threshold and dropped missing values with more than 4 consecutive days missing values. This way, we can still keep enough data, but still remove data with too many missing values. In the same way, we selected 4 days for EXACT, 1 day for heart rate, and 1 day for steps as a threshold. Regarding rescue and control medications, we did not remove any missing values and replaced them with 0 because we assumed that missing records in medications were most like that there was no medication on that day.

#### 2.3 Imputation Method

The other method to handle missing values is imputation. We evaluated 4 different imputation methods. The first one is linear imputation. The second one is deep learning-based forecasting in each patient's data. We train LSTM-based forecasting models, which take 10 days data on all features and make predictions for all features of the next day. The third one is based on the same LSTM forecasting model, but it uses FedAvg algorithm among patients. The last one is also based on the same LSTM forecasting model, but it uses our FL algorithm (Daisy-chain algorithm with clustering and curriculum learning) among patients. Figure 5 to 9 shows sample data after each imputation method is applied, and table 1 shows the comparison of the model performance. We found all

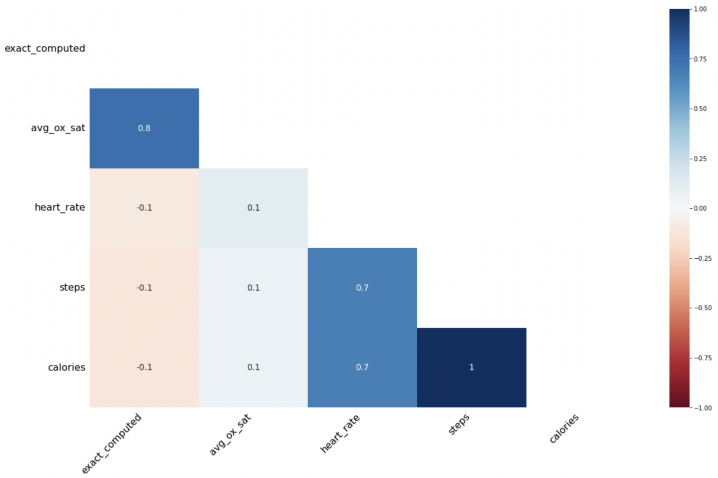


**Fig. 2.** Heatmap of missing values

results produced comparable results and did not see significant differences. This is mostly because we already dropped long consecutive missing values. Therefore, we used linear imputation for the rest of the experiments for computational efficiency.

**Table 1.** Imputation method comparison

| Imputation Method | F1 | Average Precision |
|---|---|---|
| Linear interpolation | 0.14 ±0.04 | 0.23±0.06 |
| LSTM forecasting (Local training) | 0.08 ±0.04 | 0.19±0.06 |
| LSTM forecasting (FedAvg) | 0.14 ±0.11 | 0.22±0.09 |
| LSTM forecasting (Our model) | 0.12 ±0.03 | 0.19±0.05 |

### 3 Early detection

It is also worth noting that this heterogeneity shows the potential of early detection of AECOPD because we might be able to catch a warning sign of anomalies starting from 30 days before an exacerbation event regarding survey and medication data, and 17 days before an exacerbation event regarding Fitbit data. Therefore, we conducted an experiment to see whether a deep learning model can detect AECOPD earlier. Figure 10 shows F1 score based on the test dataset for AECODP prediction 0-30 days prior to an exacerbation event. Figure 10 indicates that there is no consistent trend except that a model achieved the

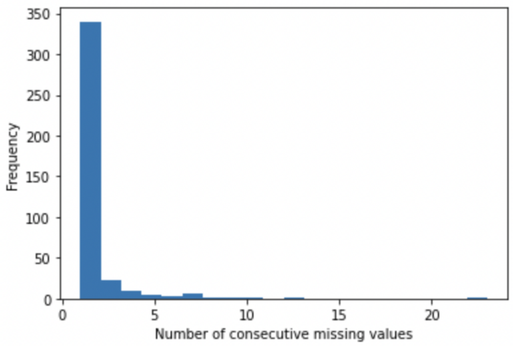


**Fig. 3.** Consecutive missing values of oxygen saturation

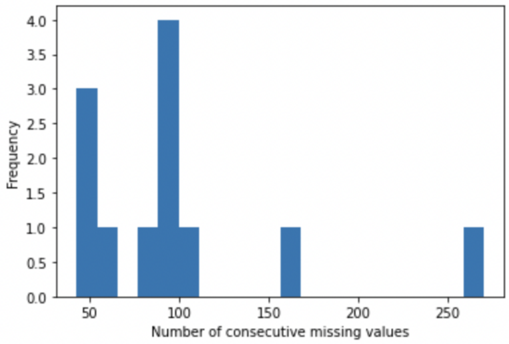


**Fig. 4.** Consecutive missing values of oxygen saturation

highest F1 score when it predicts from one day before AECOPD. This could be explained by that we might need more data to detect anomalies from earlier days since the sign of anomalies is more subtle.

### 4 Anomaly Detection Framework

#### 4.1 Weakly Supervised Learning

We performed a modified version of Label-Efficient Interactive Time-Series Anomaly Detection (LEIAD) [6]. LEIAD first creates a label function to predict labels for unlabelled data using an ensemble of unsupervised models. In addition to this, we added an ensemble process of multi-variate time series features. We made anomaly labels in cases when unsupervised methods based on more than one feature detect anomalies. Thereafter, we iterated the process of active learning

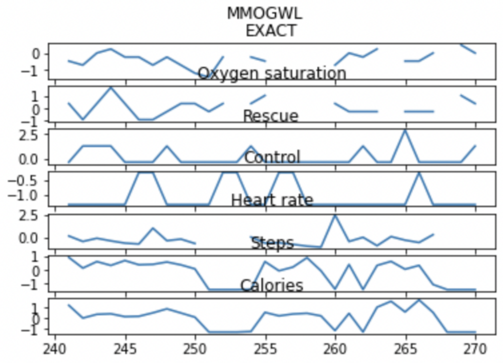


**Fig. 5.** Sample of Data (without imputation)

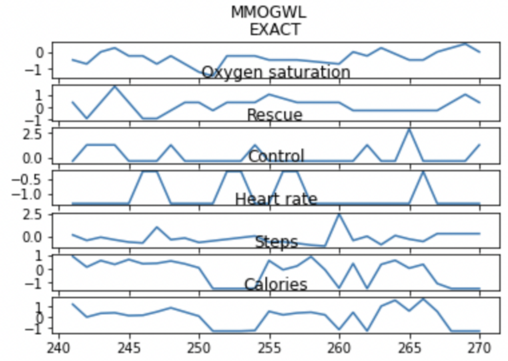


**Fig. 6.** Sample of Data (with linear interpolation)

where domain experts annotated labels, updated label functions, and trained the model. We set 5 label annotations as a minimum unit and tested the impact. Another improvement is that we use deep learning-based feature extractors, which is LSTM, instead of feature engineering such as moving average in LEIAD. This is because LSTM can do feature extractions through a learning process and this is especially useful when data has high dimensional (in our case, we have multi-variate time series data, which is high dimensional).

#### 4.2 Federated Learning

We further applied Federated Learning (FL) to share common knowledge among patients without sharing raw data. FL is a relatively new machine learning paradigm where multiple clients (e.g., smartphones or hospitals) train machine learning models collaboratively while keeping their raw data private. The most

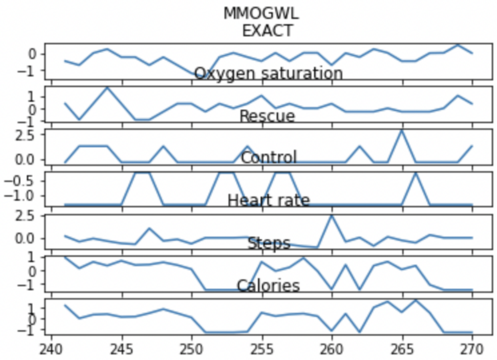


**Fig. 7.** Sample of Data (with local LSTM)

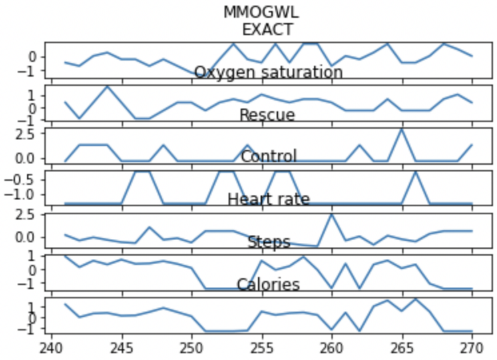


**Fig. 8.** Sample of Data (with fedavg LSTM)

common FL approach is FedAvg where each local client sends model parameters to a central server and the server sends back the average model parameters to each local client. Applying FedAvg is problematic for our data because of heterogeneity and limited data. With data heterogeneity among each user, the optimal model for each local user is different and a single global model aggregating all local models could be far from each local optimal model. In addition, since each local user lacks data in our scenario, each local user could train poor models. Aggregating local poor models is known not to work well. In order to address these challenges, we developed a novel FL approach where we combine FedDC, clustering, and curriculum learning. FedDC is our baseline model, which is recent work for small datasets. FedDC uses daisy-chain, where a central server receives one local model, and sends it to another client, the local client trains the model and sends it back to the server. This way, a model trains with all local data sequentially. FedDC could still suffer from data heterogeneity, therefore,

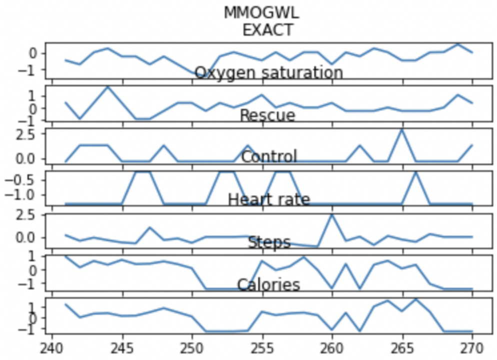


**Fig. 9.** Sample of Data (with feddc LSTM)

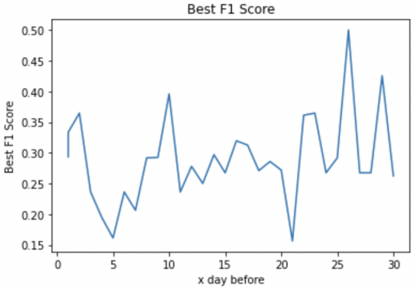


**Fig. 10.** Best F1 Score based on various timing of early detection

we apply clustering first. Instead of using a common distance measure of model parameters such as cosine similarity, we used model loss in order to measure distance based on data more directly. We first randomly select one client, train a model, send it to all other clients, compute model loss based on the locally trained model, and make a cluster if a model loss is smaller than a threshold. We iterate this process until all clients belong to some clusters. In addition to this improvement, we hypothesized that an order of daisy-chain could be crucial, so we applied curriculum learning, which is a machine learning technique where a model is trained with easy data/task first and then gradually with more difficult data/task in the same way as a human learns to master a new skill. We ordered clients within a cluster based on a model loss so that we could train from easier clients.

[6] Hong Guo, Yujing Wang, Jieyu Zhang, Zhengjie Lin, Yunhai Tong, Lei Yang, Luoxing Xiong, and Congrui Huang. Label-efficient interactive time-series anomaly detection. *arXiv preprint arXiv:2212.14621*, 2022.
